# Investigation of the Impact of Clonal Hematopoiesis on Severity and Pathophysiology of COVID-19 in Rhesus Macaques

**DOI:** 10.1101/2023.01.01.522064

**Authors:** Tae-Hoon Shin, Yifan Zhou, Byung-Chul Lee, So Gun Hong, Shayne F. Andrew, Barbara J. Flynn, Matthew Gagne, John-Paul M. Todd, Ian N. Moore, Anthony Cook, Mark G. Lewis, Kathryn E. Foulds, Robert A. Seder, Daniel C. Douek, Mario Roederer, Cynthia E. Dunbar

## Abstract

Clinical manifestations of COVID-19 vary widely, ranging from asymptomatic to severe respiratory failure with profound inflammation. Although risk factors for severe illness have been identified, definitive determinants remain elusive. Clonal hematopoiesis (CH), the expansion of hematopoietic stem and progenitor cells bearing acquired somatic mutations, is associated with advanced age and hyperinflammation. Given the similar age range and hyperinflammatory phenotype between frequent CH and severe COVID-19, CH could impact the risk of severe COVID-19. Human cohort studies have attempted to prove this relationship, but conclusions are conflicting. Rhesus macaques (RMs) are being utilized to test vaccines and therapeutics for COVID-19. However, RMs, even other species, have not yet been reported to develop late inflammatory COVID-19 disease. Here, RMs with either spontaneous DNMT3A or engineered TET2 CH along with similarly transplanted and conditioned controls were infected with SARS-CoV-2 and monitored until 12 days post-inoculation (dpi). Although no significant differences in clinical symptoms and blood counts were noted, an aged animal with natural *DNMT3A* CH died on 10 dpi. CH macaques showed evidence of sustained local inflammatory responses compared to controls. Interestingly, viral loads in respiratory tracts were higher at every timepoint in the CH group. Lung sections from euthanasia showed evidence of mild inflammation in all animals, while viral antigen was more frequently detected in the lung tissues of CH macaques even at the time of autopsy. Despite the lack of striking inflammation and serious illness, our findings suggest potential pathophysiological differences in RMs with or without CH upon SARS-CoV-2 infection.

**Highlights:**

- No evidence of association between CH and COVID-19 clinical severity in macaques.
- The presence of CH is associated with prolonged local inflammatory responses in COVID-19.
- SARS-CoV-2 persists longer in respiratory tracts of macaques with CH following infection.

## INTRODUCTION

Despite the progress in decreasing morbidity and mortality due to COVID-19 with the advent of antivirals and vaccines, there remain unresolved issues in disease pathophysiology and treatment. While most patients show asymptomatic or self-limited mild illness, a considerable number of patients progress to life-threatening respiratory failure often accompanied by serious complications, including thrombosis, coagulopathy, and cardiovascular disease (CVD). This deterioration often occurs during the second week following infection, with evidence implicating a hyperinflammatory state as associated with poor outcomes [1,2,3]. Extensive studies to date have uncovered several risk factors for severe COVID-19 disease, most strikingly age, as well as gender, ethnicity, underlying genetic variation in immune response pathways, and comorbidities such as diabetes, obesity, and hypertension [4,5,6]. However, even within high-risk groups such as the elderly, outcomes are very heterogeneous and cannot be fully explained by known factors predicting severe illness.

Acquired somatic mutations in hematopoietic stem and progenitor cells (HSPCs) are associated with an *in vivo* competitive advantage and certain human health outcomes including myeloid malignancies. This syndrome is termed clonal hematopoiesis (CH) and greatly increases with advanced age, in a pattern that mirrors the association between age and COVID-19 severity. Of note, loss-of-function (LOF) mutations in certain genes such as the epigenetic regulators *DNMT3A* and *TET2* are the most frequent CH mutations and have been linked to a distinct myeloid cell phenotype associated with hypersecretion of inflammatory cytokines and an increased risk of CVD [7,8,9]. Thus, we and others have hypothesized that the presence of CH could influence the risk of severe COVID-19 disease. Although several human cohort studies have examined this relationship, results to date are conflicting and difficult to interpret due to insufficient sample size, inconsistent sequencing platforms, inclusion of cancer patients with competing risks for CH, and limited availability of information on other potential risk factors [10,11,12,13,14]. This potential relationship has not yet been studied in a preclinical model.

Various animal models for COVID-19 have been developed, although no one model closely mimics the course or severity of human disease (reviewed in [15] and [16]). Rhesus macaques (RMs) have been established as the most relevant model for SARS-CoV-2 infection and are being widely utilized for preclinical development of therapeutics and vaccines [17,18]. However, preclinical models that recapitulate the pathophysiology of severe or fatal human disease with late hyperinflammation do not yet exist. Macaques experimentally infected with SARS-CoV-2 do not succumb to the infection or develop serious hyperinflammatory clinical manifestations.

We previously identified naturally occurring CH in aged RMs carrying a spectrum of mutations and an incidence similar to human CH [19]. In addition, given the lack of availability of naturally aged macaques, we created a CRISPR/Cas genetically engineered CH model in younger adult macaques that reproduces the expansion of clones with TET2 LOF and hyperinflammation [19]. In this pilot study, we utilized our macaque models to ask whether disease severity and biologic characteristics following SARS-CoV-2 inoculation was enhanced in animals with either naturally occurring CH expanding post-autologous transplantation or engineered CH mutations in the *DNMT3A* or *TET2* genes introduced via *ex vivo* editing followed by autologous transplantation, in comparison to similarly transplanted and conditioned middleaged adult animals without CH mutations.

## MATERIALS AND METHODS

Additional details are available in Supplementary Materials.

### Animals and SARS-CoV-2 challenge

All procedures were approved by the Animal Care and Use Committee of the National Heart, Lung, and Blood Institute, Vaccine Research Center and Bioqual (Rockville, MD).

Macaques with either natural (n=1) or engineered CH (n=2) along with autologously-transplanted conditioned controls (n=3) were inoculated intra-bronchially with 8 x 10^5^ PFU total of SARS-CoV-2 (USA-W1/2020 strain) and monitored until 12 days post-inoculation (dpi).

### Histopathology

Lung tissue samples were fixed, processed, and embedded in paraffin. Sections from three different lobes were stained with hematoxylin and eosin or immunohistochemical reagents. Scoring criteria for inflammation and viral detection are detailed in Supplementary Methods.

### Viral loads

Subgenomic RNA (sgRNA) for SARS-CoV-2 envelope and nucleocapsid were quantified in nasal swab specimens and bronchoalveolar lavage fluid (BALF) via PCR as previously described [17].

### Cytokine quantification

Cytokine/chemokine levels in serum and BALF were analyzed using the MILLIPLEX MAP Nonhuman Primate Cytokine Magnetic Beads Panel (Millipore Sigma, Burlington, MA) and MAGPIX with Bio-Plex Manager™ MP software (Bio-Rad, Hercules, CA).

### Clonal tracking

Changes in variant allele frequency (VAF) of CH mutations following infection were analyzed in granulocytes and cellular components of BALF via targeted deep sequencing and custom pipeline as reported [19].

## RESULTS AND DISCUSSION

Previously described RMs with spontaneous *DNMT3A* CH expanding post-autologous transplantation (RQ859) [19,20] or CRISPR-engineered *TET2* CH (ZL39 and ZH63) [19], along with control non-CH animals receiving autologous transplantation following the same total body irradiation conditioning protocol (n=3) were inoculated with SARS-CoV-2 and monitored for 12 days (Figure 1A). All animals manifested only mild signs of clinical illness, without differences between the two groups, and no evidence of significant pulmonary inflammation on repeated chest radiographs (Figure S1). However, the natural aged CH animal RQ859 died suddenly on 10 dpi of unclear causes, however, the event may have been due to unexpectedly severe baseline anemia present at viral challenge (Figure S2), but an effect of SARS-CoV-2 infection cannot be excluded. RMs with CH showed a slower tendency in body weight recovery compared to controls, even excluding RQ859 (given the possibility of another cause for mortality in this animal), although the differences were not statistically significant due to the sample size. No remarkable differences were observed in body temperature or respiratory and heart rates between the groups (Figure 1B).

**Figure 1.**
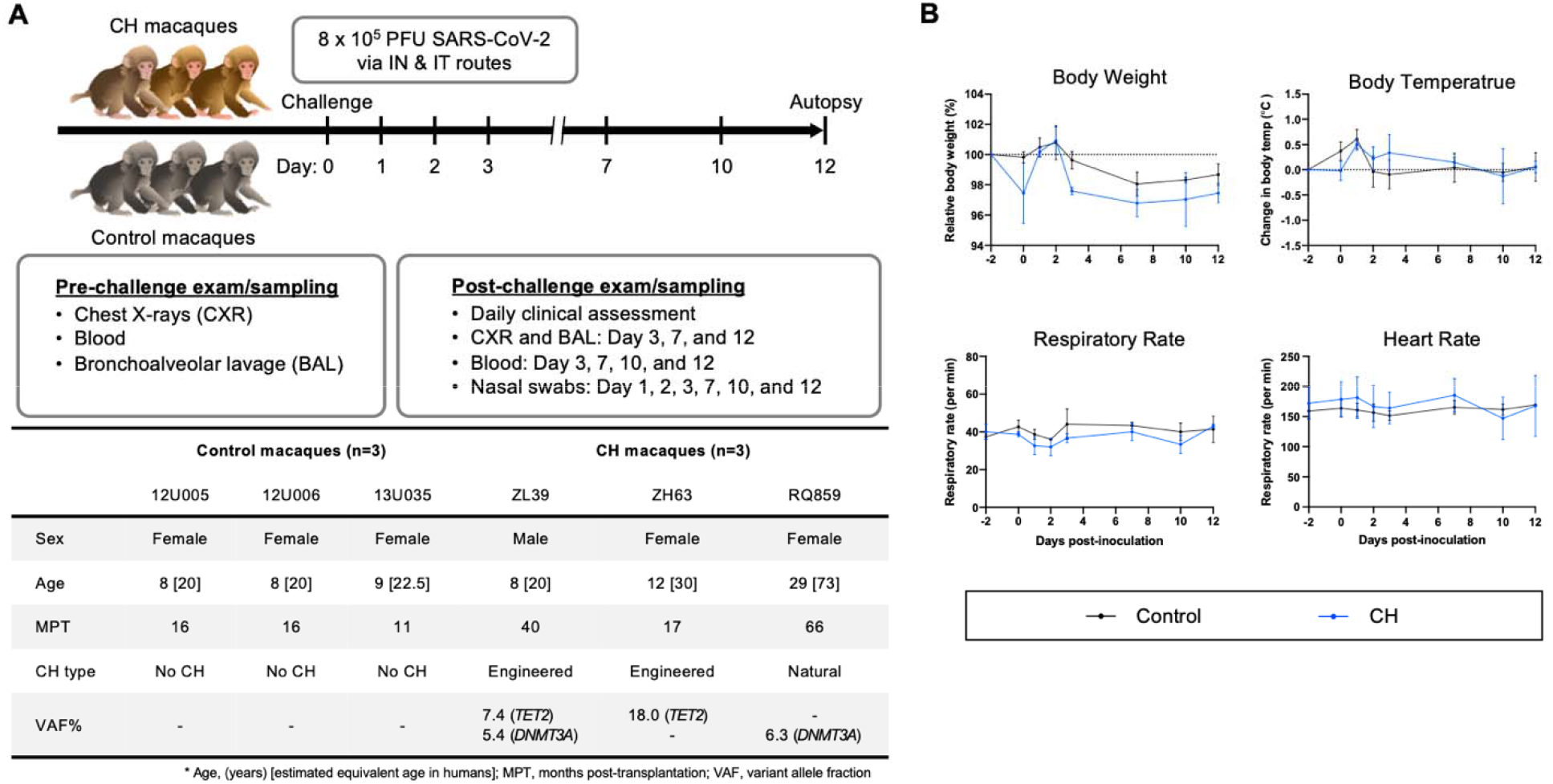
SARS-CoV-2 challenge in a RM model of Clonal Hematopoiesis. (A) Schematic workflow of the experimental procedures, including SARS-CoV-2 inoculation, clinical and radiographic examinations, and specimen collection. Additional information regarding macaques enrolled in this study at the time of the SARS-CoV-2 inoculation are summarized in the table. The number in brackets in the age row gives the estimated equivalent age in humans. In control animals, absence of CH somatic mutations was verified by error-corrected ultra-deep exome sequencing of a 56 gene panel associated with myeloid neoplasms [19]. All animals receiving myeloablative total body irradiation conditioning prior to autologous transplantation. All animals had normal hematopoiesis as assessed by blood counts and marrow morphology at study entry, however animal RQ859 was unexpectedly found to be severely anemic on baseline sampling taken at the time of inoculation. IN, intranasal; IT, intratracheal. (C) Relative body weight and body temperature changes following inoculation and changes in respiratory and heart rates between CH and control group.

To evaluate post-challenge inflammatory responses in the presence or absence of CH, sera and BALF cytokine/chemokine concentrations were measured serially. In BALF, mean concentrations of MCP-1, IL-6, IL-8 and MIP-1β were consistently higher in CH macaques compared to controls (Figure 2A), whereas no constant patterns in serum cytokine/chemokine levels were observed between the two groups, except for somewhat higher IL-6 concentration in CH animals until 12 dpi (Figure S3). These results are consistent with a study demonstrating sustained local inflammatory responses in older versus younger RMs, despite similar disease outcomes [21], suggesting the possibility of CH-specific divergence of immune responses against SARS-CoV-2 even at similar ages. The prior study did not screen for CH in the older macaques.

**Figure 2.**
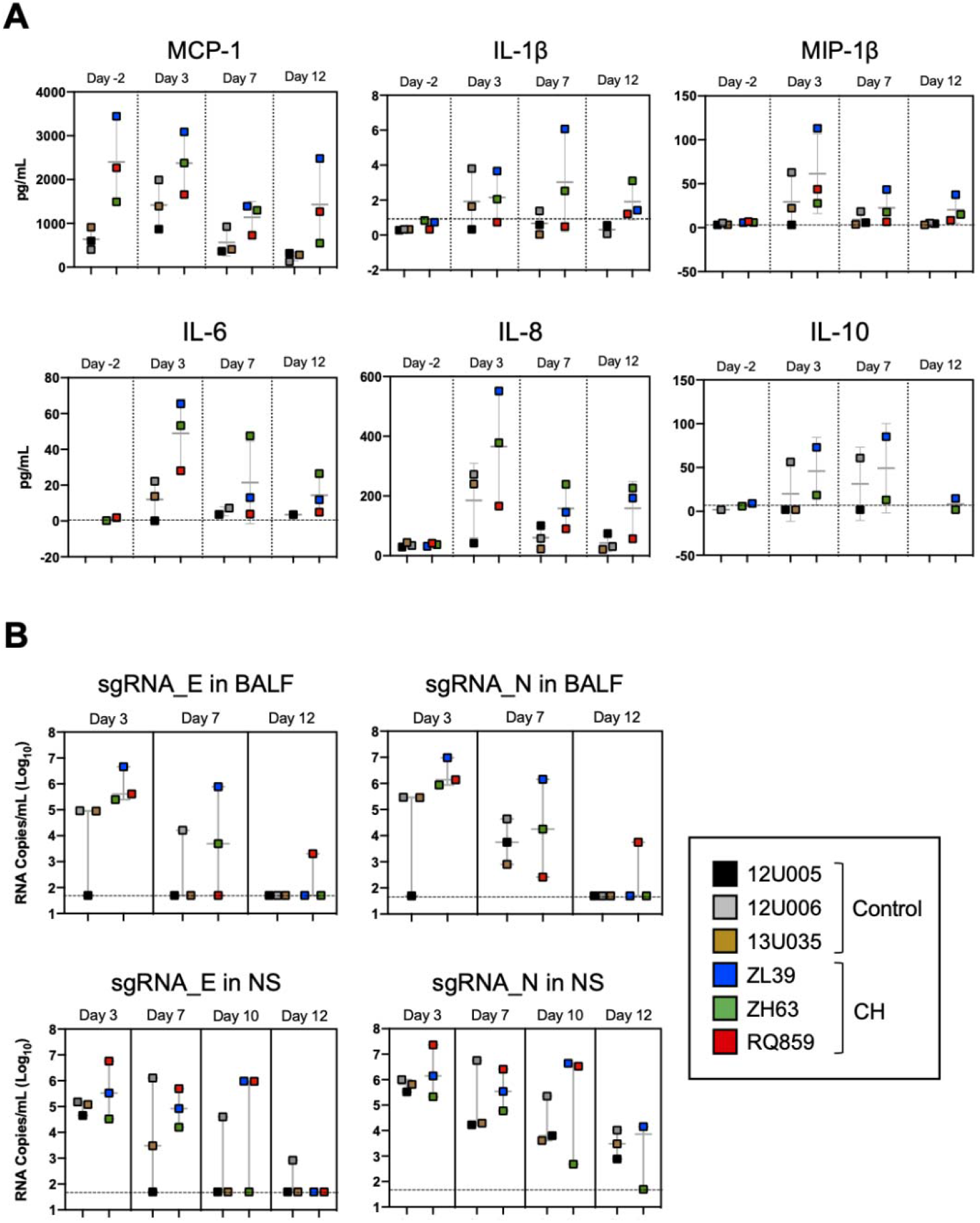
Post-challenge inflammatory response and viral load in macaques with CH. (A) Post-challenge BALF cytokine/chemokines levels in CH macaques versus controls. Box plots depicting the results of immunoassays for rhesus macaque MCP-1, IL-1β, MIP-1β, IL-6, IL-8 and IL-10. (B) Assessment of viral replication in BALF and nasal swab specimens by SARS-CoV-2 envelope (E) and nucleocapsid (N) sgRNA quantification. Each colored symbol indicates individual macaques and the dashed horizontal lines show the limit of detection for the assay.

Given that IL-6 is hypothesized to be a major factor contributing to the vicious cycle between hyperinflammation and clonal expansion in *TET2* and *DNMT3A* CH [22,23,24], we asked whether the elevated levels of lung IL-6 in CH macaques resulted from or contributed to local lung expansion or persistence of CH mutant cells following SARS-CoV-2 infection. Targeted deep sequencing measured CH mutation VAFs in both BALF cells and granulocytes isolated from PB and BM. No significant changes were observed in mutation VAFs in PB/BM granulocytes following infection. In BALF, VAFs shifted in both directions following infection (Figure S4), possibly due to changes in neutrophil to lymphocyte ratio with infection, since VAFs are usually lower in lymphoid cells in the presence of CH [19].

We measured levels of sgRNAs encoding SARS-CoV-2 envelope and nucleocapsid in nasal swabs and BALFs collected at 3, 7, 10 and/or 12 dpi to assess viral replication in the upper and lower airways. Interestingly, the median copy numbers of both sgRNAs were higher at every timepoint in the CH group compared with controls, with more substantial differences in BALF (Figure 2B), suggesting that CH might impact on the extent of viral replication in both the upper and lower respiratory tracts. Histopathologic analysis on the lung sections from euthanasia at 10 or 12 dpi found evidence for mild inflammation in all animals (Figure 3A). Confirming the viral load measurements, viral antigen was immunohistochemically detected in the lung tissues of all three CH animals but only one of three controls (Figure 3B). No remarkable differences were found in other tissues such as lymph nodes and spleen, and we did not find any atherosclerotic changes in the hearts of the CH animals. Although the relationship between viral load and COVID-19 severity has not yet been established with certainty [25,26,27], our findings, together with a recent study showing a higher level and longer period of viral RNA detection in elderly macaques [28], are relevant to considering the relationship between aging, CH and COVID-19.

**Figure 3.**
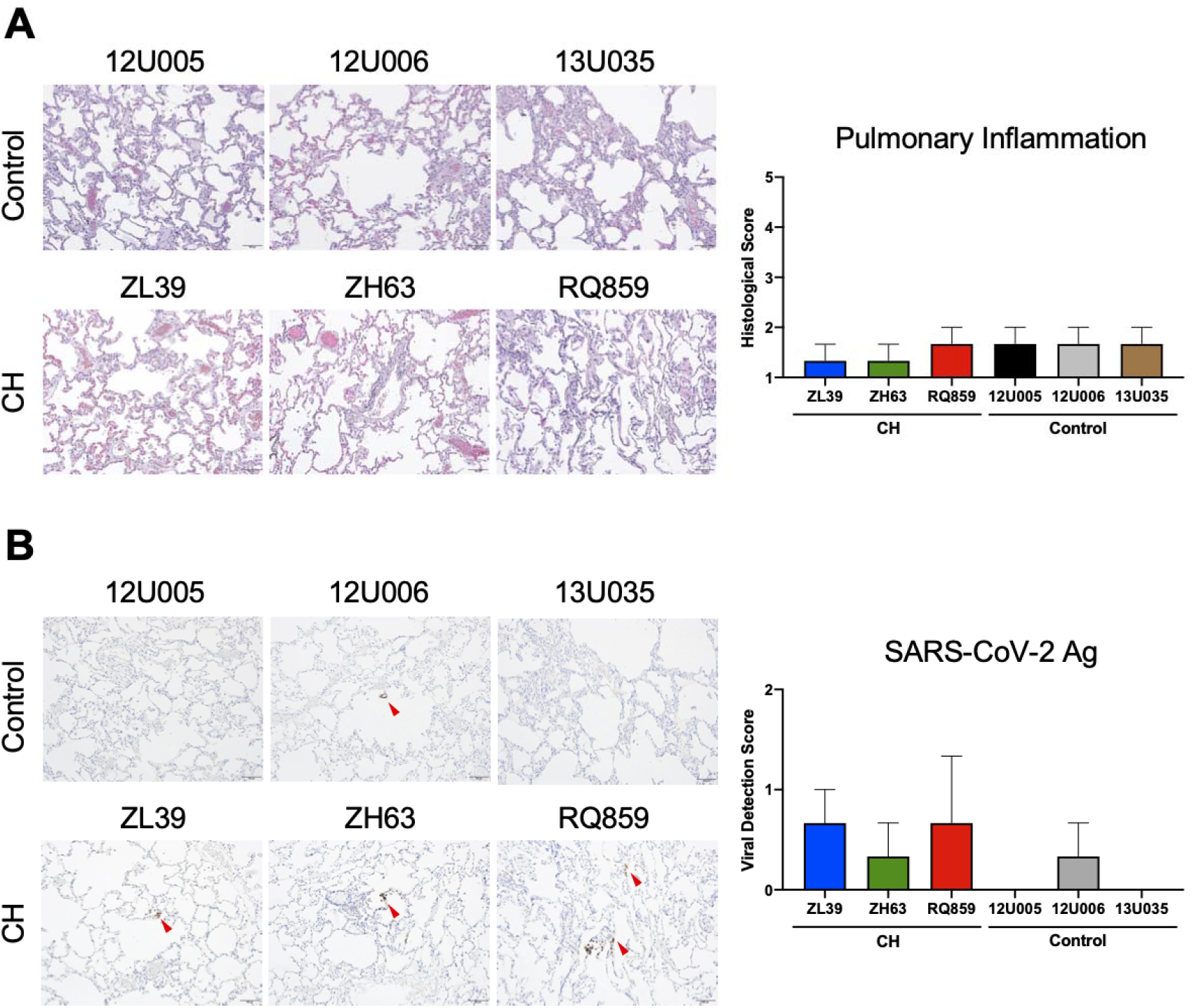
Histopathologic and immunohistochemical analyses of the lung tissue. To assess the degree of pulmonary inflammation and virus survival, the lung tissues of RMs with or without CH isolated at 12 dpi were stained with H&E (A) and an antibody specific for SARS-CoV-2 Ag (B), respectively. Representative microscopic images are shown on the left side of each panel (scale bar = 100 μm), and histopathologic and viral detection scores were calculated based on the defined criteria (details in Supplementary Methods) and graphed on the right. Red arrowheads in panel B indicate the detected viral antigens in the lung tissues.

In summary, our nonhuman primate model of CH demonstrates its possible utility as a preclinical tool for investigating the relationship between CH and other diseases such as COVID-19. Consistent with our recently completed study in large human cohorts [14], we found no evidence for a direct correlation between CH and severity of clinical COVID-19 in RMs, although data from this small cohort does provide interesting preliminary evidence for potential pathophysiological differences in macaques with or without CH upon SARS-CoV-2 infection, particularly regarding the level of inflammatory cytokines/chemokines in the lung, and the degree and persistence of virus in the upper and lower respiratory tracts. The creation of CH in macaques is straightforward, and this model may be useful for hypothesis testing in the future.

## ACKNOWLEDGEMENTS

We thank the Bioqual and NHLBI Primate Program, Nathaniel Linde, Allen Krouse, Theresa Engels, Justin Golomb, and the veterinary staff for care of the macaques.

## AUTHOR CONTRIBUTIONS

T-H.S. and C.E.D conceptualized the study. C.E.D., K.E.F., R.A.S, D.C.D., and M.R. designed and supervised the study. T-H.S., Y.Z., B-C.L., S.F.A., B.J.F., M.G., I.G.M., and A.C. performed experiments and analyzed data. S.G.H., J-P.M.T., and M.G.L. organized and conducted the animal care, transfer, and infection. T-H.S. and C.E.D. wrote the manuscript with input from all coauthors.

