## Supplementary Materials for "Investigation of the Impact of Clonal Hematopoiesis on Severity and Pathophysiology of COVID-19 in Rhesus Macaques"

**SUPPLEMENTARY METHODS**

***Animals and SARS-CoV-2 challenge***

All procedures were approved by the Animal Care and Use Committee of the National Heart, Lung, and Blood Institute, Vaccine Research Center and Bioqual (Rockville, MD). Infection experiments were conducted in animal biosafety level 3 facility at Bioqual.

One aged macaque with natural DNMT3A CH and two with engineered TET2 CH, along with three control animals similarly transplanted and conditioned, were inoculated with SARS-CoV-2 (USA-W1/2020 strain). Animals were challenged with 8 x 10^5^ plaque-forming units (PFU) total of SARS-CoV-2 via intranasal and intratracheal routes as described previously [1] and monitored for 12 days. Strategies for exams and specimen collections are detailed in Figure 1A.

***Clinical assessment***

Cage-side clinical assessment was performed daily with a detailed scoring sheet. Animals were anesthetized for chest X-ray, complete blood count, blood and bronchoalveolar lavage fluid (BALF) draws on 2 days pre- and 3, 7, 10, and 12 days post-inoculation (dpi) and for nasal swabs on days 1, 2, 3, 7, 10, and 12. During anesthesia, vital indices such as body weight, temperature, and respiratory and heart rates were monitored.

***Viral loads***

To quantify SARS-CoV-2 replicating in respiratory tract, polymerase chain reaction (PCR) was performed to detect subgenomic RNAs (sgRNAs) of both envelope (E) and nucleocapsid (N) as described previously [1,2]. Briefly, total RNA was extracted from nasal swabs and BALF using RNAzol BD column kit (Molecular Research Center, Cincinnati, OH) according to the manufacturer’s instruction. PCR reactions were conducted using TaqMan Fast Virus 1-Step Master Mix (Applied Biosystems, Waltham, MA), gene-specific probes, forward primer in the 5’ leader region, and reverse primers. Amplification of sgRNAs was calculated using the QuantStudio 6 Pro Real-Time PCR System (Applied Biosystems), with a lower detection limit of 50 copies per reaction.

***Cytokine quantification***

Serum was separated from cellular components of each blood sample by centrifugation at 4°C. BALs were concentrated 10X prior to cytokine quantification. Concentrations of major cytokines and chemokines in serum and BAL were measured using MILLIPLEX MAP Nonhuman Primate Cytokine Magnetic Beads Panel (Millipore Sigma) according to the manufacturer’s instructions. Luminex data were analyzed using MAGPIX with Bio-Plex Manager™ MP software (Bio-Rad).

***Clonal tracking***

Through density gradient centrifugation, mononuclear cells and granulocytes were isolated from peripheral blood (PB) and bone marrow (BM). Cellular components of BAL were separated from fluid after centrifugation. Genomic DNAs extracted from the granulocytes and BAL cells were amplified using gene-specific primers for mutated regions at DNMT3A (RQ859) and TET2 (ZL39 and ZH63) and ligated with unique index sequences, followed by sequencing on Illumina Miseq. The sequencing reads with mean depth of more than 300,000 were analyzed using CRISPResso (<http://crispresso.rocks>) and custom R pipelines as documented previously [3]. The remaining portion of the BAL cells were used for immunotyping of the population using flow cytometry.

***Histopathology***

Lung, spleen, and lymph node samples were collected from SARS-CoV-2-infected macaques during autopsy performed on day 12. The tissues were fixed in buffered formalin, processed with tissue processor, embedded in paraffin, and sectioned serially at 5 μm. Subsequently, the sections were stained with hematoxylin and eosin (H&E). Immunohistochemical (IHC) analysis was performed with formalin-fixed paraffin-embedded lung sections obtained from at least three different lobes per animal. As described previously [1], the slides were stained with a rabbit polyclonal SARS-CoV-2 (GeneTex, Irvine, CA) using the BOND-RX Multiplex IHC Stainer (Leica Biosystems, Wetzlar, Germany), and DAB chromogen was detected using the Bond Polymer Refine Detection Kit (Leica Biosystems). Microscopic evaluation of each staining was conducted by a board-certified veterinary pathologist. Inflammation and viral detection scores were calculated according to the following scales: 1) 0 = absent, 1 = minimal to mild, 2 = mild to moderate, 3 = moderate to severe, 4 = severe pulmonary inflammation, 2) 0 = non-detected, 1 = rare to occasional, 2 = occasional to multiple, 3 = multiple to numerous, 4 = numerous SARS-CoV-2 antigens.

***Statistical analysis***

All statistical analyses were performed using GraphPad Prism 9 (GraphPad Software, San Diego, CA). All graphs with error bars are presented as the mean ± S.E.M. Unpaired student’s t-test was applied for pairwise comparisons between two groups, and One-way ANOVAs followed by the Tukey’s post-hoc test for comparisons of multiple groups.

**SUPPLEMENTARY FIGURES**

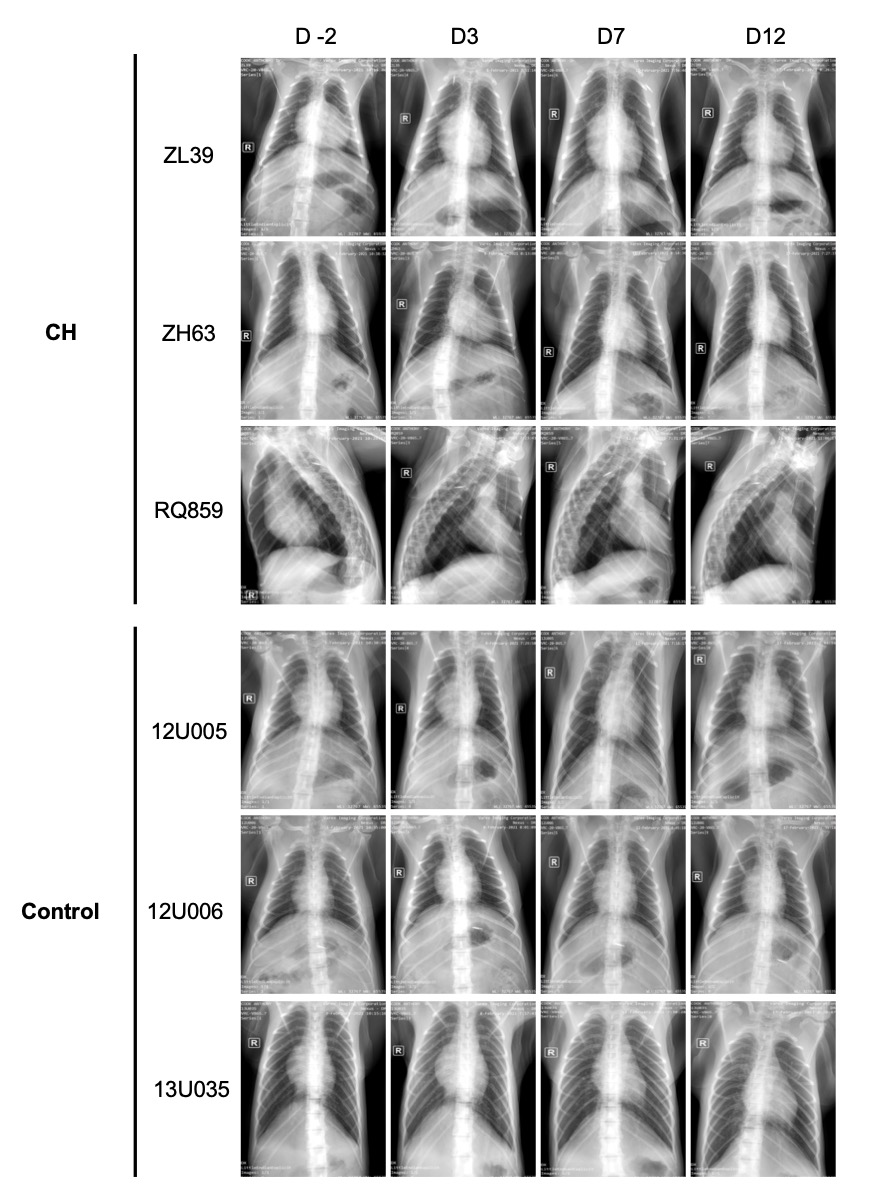

**Figure S1. Post-challenge chest radiographs in macaques with or without CH.**

Chest radiographs were taken repeatedly to assess pulmonary infiltrates upon SARS-CoV-2 inoculation. For RQ859, the image taken on 10 dpi replaces the 12 dpi due to the sudden death before autopsy.

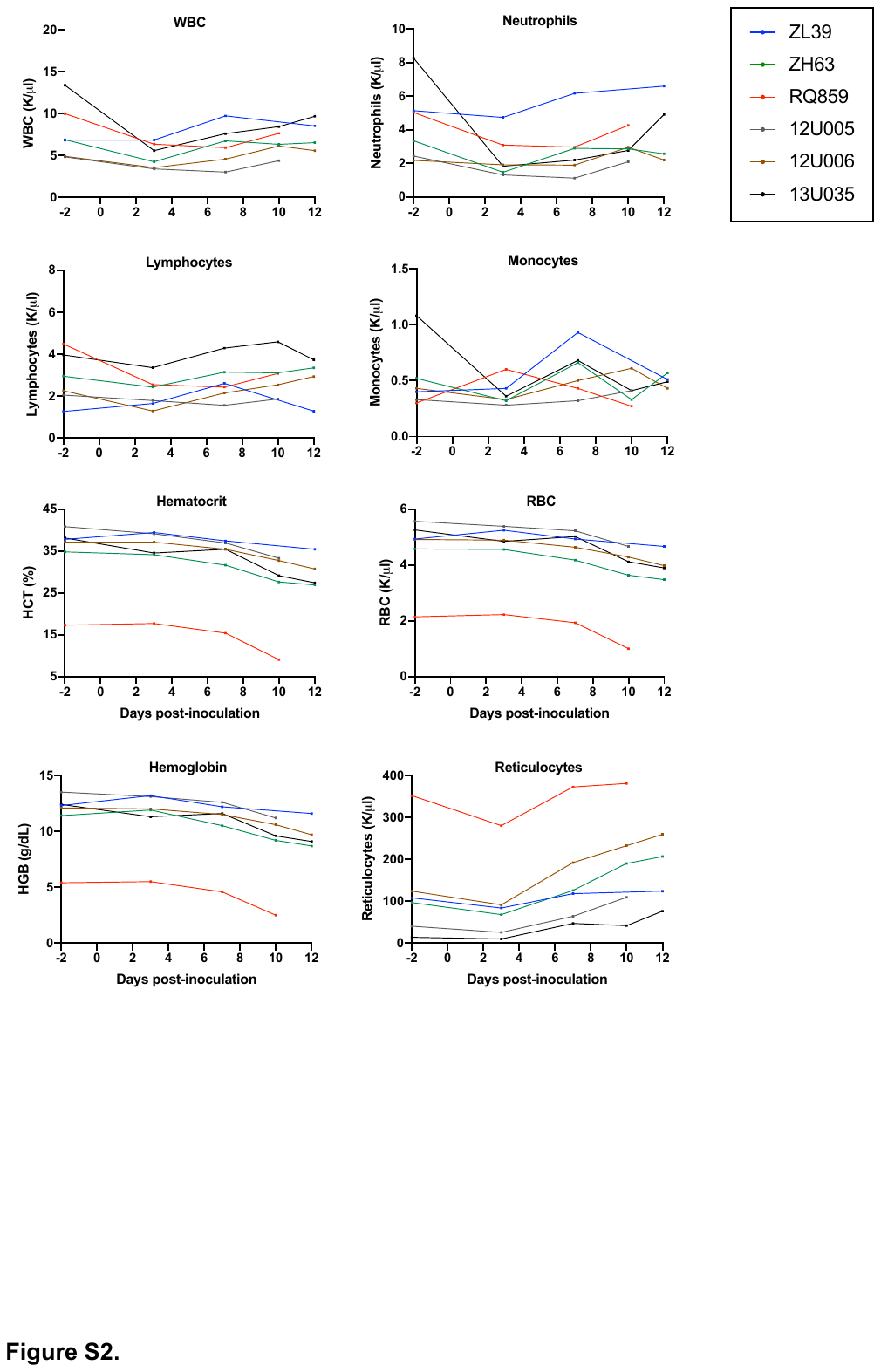

**Figure S2. Complete blood counts of macaques with or without CH after inoculation.**

Changes in the primary indices of blood counts before and after challenge were indicated for each animal. RQ859 was unexpectedly severely anemic at the time of the challenge.

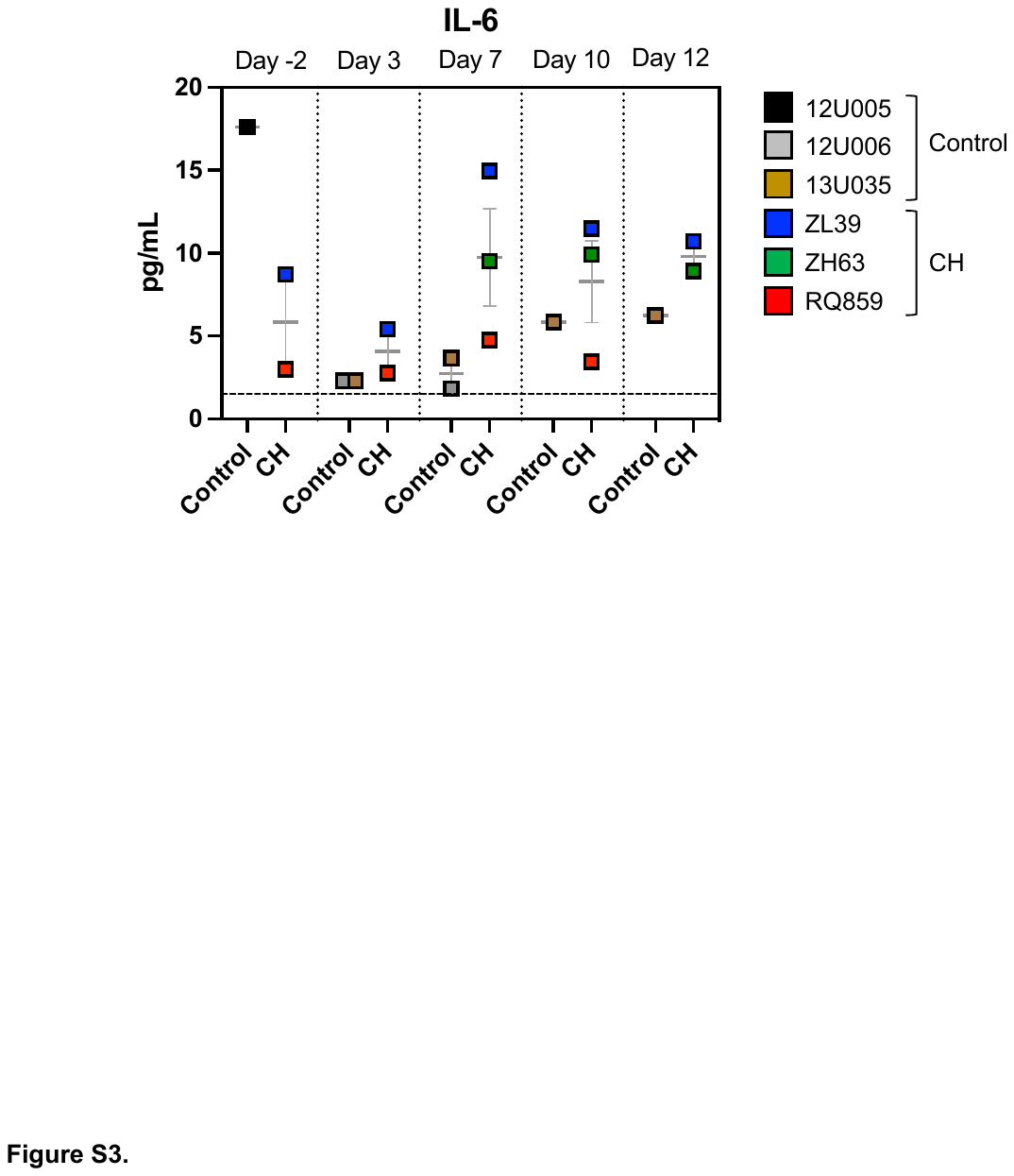

**Figure S3. Serum IL-6 levels following SARS-CoV-2 infection.**

Blood serum samples collected on baseline (-2) and 3, 7, 10, and 12 dpi were analyzed for a panel of 23 chemokines and cytokines using MILLIPLEX® MAP. Among them, the concentration change of macaque IL-6 was depicted here. Data are presented as the mean ± S.E.M. Each colored symbol indicates individual animals, and the dashed horizontal line shows the detection limit of the assay.

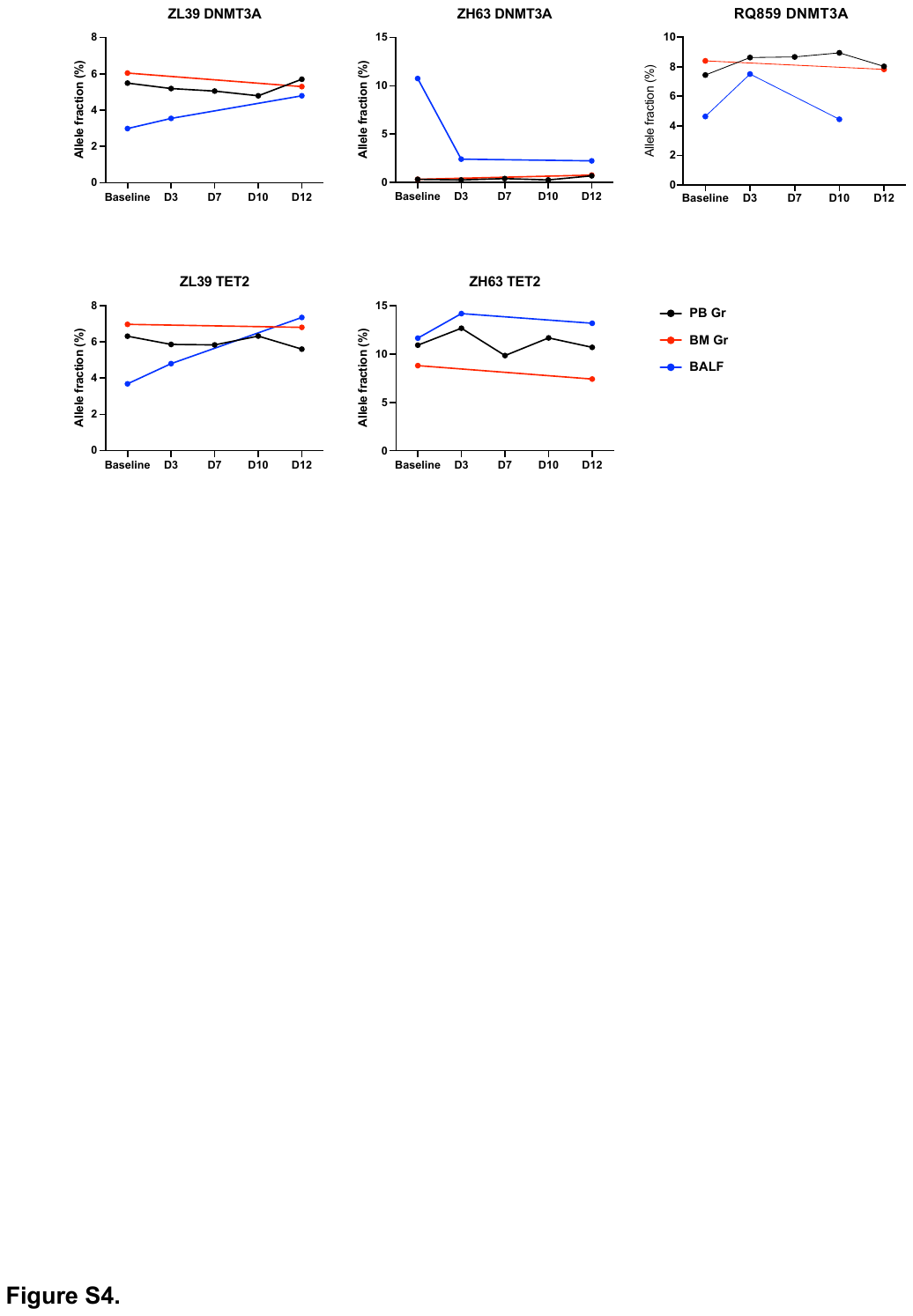

**Figure S4. Tracking of CH mutations following infection in blood and lung of CH macaques.**

Pre- and post-inoculation changes in VAFs of *DNMT3A* mutations (engineered mutations of ZL39 and ZH63 and spontaneous mutation of RQ859, respectively) and TET2 mutations (ZL39 and ZH63) were analyzed in granulocytes (Gr) from peripheral blood (PB) and bone marrow (BM) and cellular components of BALF using targeted deep sequencing.
